# An intelligent synthetic bacterium with sound-integrated ability for chronological toxicant detection, degradation, and lethality

**DOI:** 10.1101/2022.06.08.495251

**Authors:** Huan Liu, Lige Zhang, Weiwei Wang, Haiyang Hu, Ping Xu, Hongzhi Tang

**Affiliations:** State Key Laboratory of Microbial Metabolism, Joint International Research Laboratory of Metabolic and Developmental Sciences, and School of Life Sciences and Biotechnology, Shanghai Jiao Tong University, Shanghai, People’s Republic of China

**Keywords:** integrated system, autonomy, biosensor, degradation, biological safety

## Abstract

Modules, toolboxes, and systems of synthetic biology are being designed to solve environmental problems. However, weak and decentralized functional modules require complicated controls. To address this issue, we investigated an integrated system that can complete detection, degradation, and lethality, in chronological order without exogenous inducers. Biosensors were optimized by regulating expression of receptor and reporter to get higher sensitivity and output signal. Several stationary-phase promoters were selected and compared, while promoter P_*fic*_ was chosen to express the degradation enzyme. We created two concepts of lethal circuits by testing various toxic proteins, with a toxin/antitoxin circuit showing a potent lethal effect. Three modules were coupled, step-by-step. Detection, degradation, and lethality were sequentially completed, and the modules had partial attenuation compared to pre-integration, except for degradation. Our study provides a novel concept for integrating and controlling functional modules, which can accelerate the transition of synthetic biology from a concept to practical applications.

**Teaser:** We provide new ideas for integration and chronological control of multiple modules in synthetic biology.

## Introduction

Environmental pollution performs a more and more profound impact on national health and economic development, while synthetic biology has shed new light on solving this problem. Therefore, several synthetic modules, tools, and systems are being designed focusing on three main aspects: detection, degradation, and biological safety (*1*). Regarding detection, whole-cell biosensors have attracted attention because of their low cost, high selectivity, and ease of manufacturing (*2*). Nucleic acid- and protein-based biosensors are the two most common types that are able to undergo conformational alterations upon input ligand binding to regulate the expression of output signals (*3*), such as the guanidine-bound *S. acidophilus* guanidine-I riboswitch, ArsR for arsenic detection, MerR for mercury detection, and DmpR for detecting organophosphate pesticides containing phenolic groups (*4-6*). Regarding degradation, analysis of the catabolic pathways in natural strains facilitates the migration of functional genes to artificial cells that do not possess efficient or complete degradation abilities (*5, 7*). For example, an artificial consortium of three *E. coli* BL21(DE3) strains with synergistic functional modules was designed to completely degrade phenanthrene (*8*), and the pathway of engineered *E. coli* DH5α was linked with a natural pentachlorophenol degrader to mineralize hexachlorobenzene (*9*). Additionally, restraining their proliferation is one of the primary challenges for genetically modified microorganisms. There are many pioneering biocontainment strategies, including engineered prevention of self-replication, auxotrophy, synthetic gene circuits, and integrated killing systems (*10, 11*).

However, weak and decentralized functional modules for detection, degradation, and biosafety require comprehensive control conditions that hinder the ability of synthetic biology to solve environmental problems. First, the biosensor needs a stable contaminant concentration to produce a reliable output signal, which limits the time for strains to begin degradation, but favors high degradation rates like high-density fermentation (*12*). It is then necessary to kill the engineered cells after completing the degradation of target compounds, but most strategies depend on exogenous inducers or physical conditions (*11*). Finally, it is quite important to keep each module of the combination robust. Studies have attempted to achieve these goals. For example, a 9 kb naphthalene-degrading gene *nahAD* was cloned to *Acinetobacter* ADPWH_lux capable of responding to salicylate (*13*). An efficient Hg^2+^ adsorption strain with a biocontainment system was designed, and it achieved Hg^2+^ adsorption efficiency >95% with an escape rate <10^−9^. However, there is still an urgent lack of a synthetic biology system capable of autonomously and efficiently performing multiple integrated functions in chronological order without exogenous chemical inducers (*14*).

To meet these challenges, we selected salicylic acid (SA), a typical pollutant in industrial wastewater, as the target compound. We assembled a three-module engineered strain that could first efficiently detect and degrade SA, after which the suicide module autonomously activated. All tasks were completed sequentially by the engineered strain without any intervention. We selected several promoters and ribosome binding sites to regulate the receptor’s density and the reporter’s intensity, optimizing key biosensor parameters such as sensitivity and dynamic range. To turn on the degradation module, various stationary-phase promoters were tested. To prevent the escaping of the engineered strains when SA was completely consumed, two lethal circuits were compared, and the better one with higher lethal efficiency was integrated. We assembled these modules, step-by-step, and tested their functions. Thus, our study sheds new light on the functional normalization and timing control of synthetic biology modules used to treat environmental pollutants.

## Results

### Construction and characterization of biosensors for salicylic acid

NahR, a LysR-type transcriptional activator of the *nah* and *sal* promoters, responds to salicylate (*15*) and can be constructed as a biosensor to conveniently detect SA concentrations at a low cost. A wild-type biosensor (WT) was constructed using mRFP as the reporter (Fig. 1A), but its low response, narrow dynamic range, and detection range limited its application (Fig. 1F–1H and Table 1). Therefore, this sensor was optimized in two ways, regulating receptor density and reporter intensity. Promoters and ribosome binding sites (RBS) with different intensities were selected from the iGEM registry (https://old.igem.org/Registry) to determine the optimal combination (Fig. 1A). J231XX-B003X represents the biosensor optimized by each promoter (J231XX) and RBS (B003X) (Table 1 and Fig. 1). To avoid the effect of SA on strain growth, the growth curve under gradient concentrations of SA was plotted (Fig. 1B), while 0.001–5,000 μM SA did not inhibit the cell growth. In the pre-selection at a 96-well plate, only three combinations showed obvious improvements (Fig. 1C–1E). Compared with WT, their detection limits decreased to 0.1 mM and the response value increased from approximately 50 to a maximum of 600 at 8 h. However, there were also several disadvantages, with leakage on J23105-B0030 at 8 and 10 h, and FI/OD_600 nm_ reached saturation at a lower concentration.

**Figure 1.**
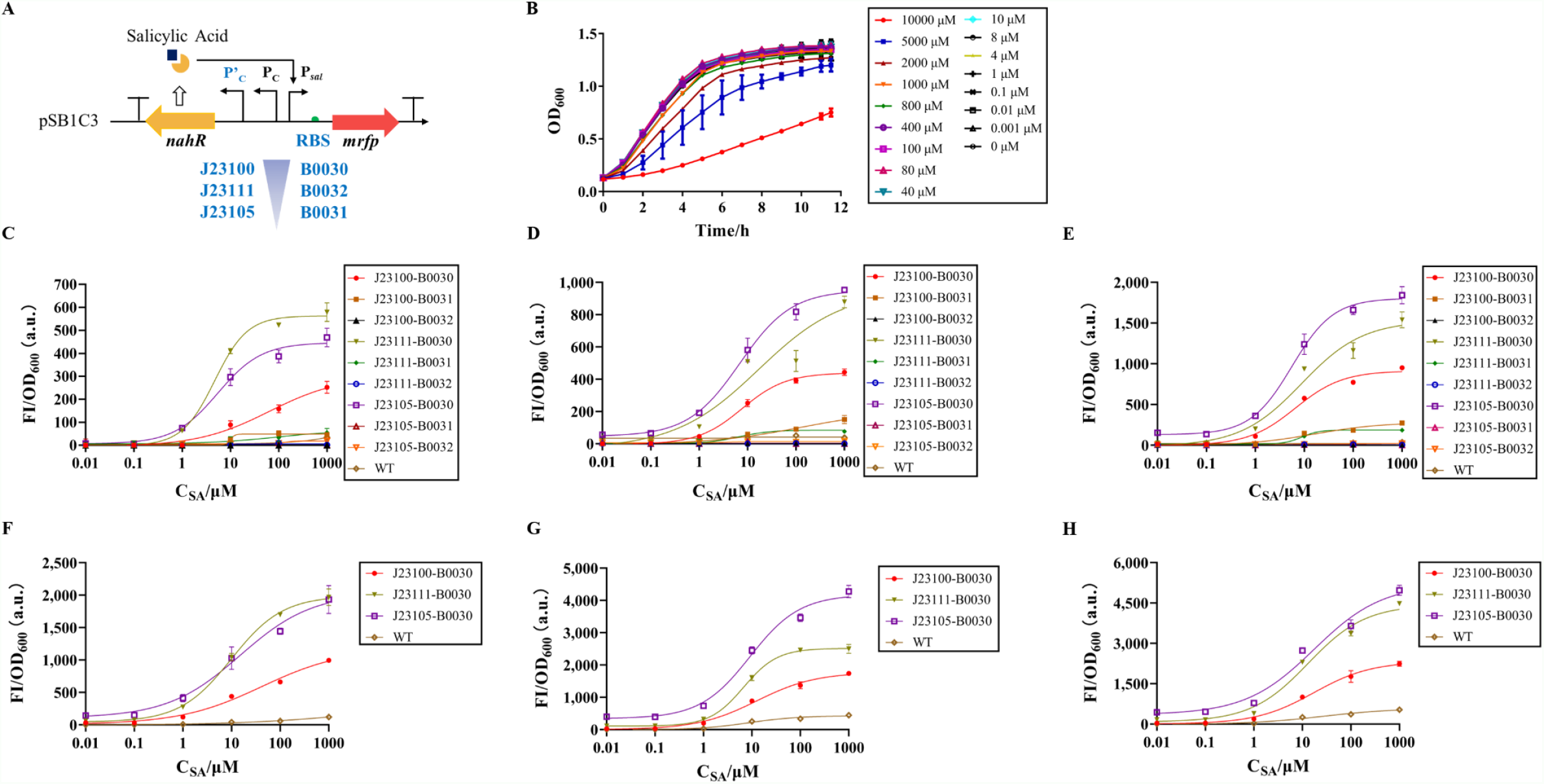
The principle and optimization of the biosensor for SA. (**A**), NahR is a LysR-type activator of *nah* and *sal* promoters responding to SA. We used promoters of different strengths to regulate the density of the receptor, and ribosome binding sites (RBS) of different strengths to adjust the reporter expression. J231XX and B003X are registration numbers of promoters and RBS in the iGEM registry. (**B**), Growth curves of *E. coli* Top10 at different concentrations of SA. (**C–E**), Dose-Response curves of the biosensor with different optimized combinations in 96-well plate, C–E are the results for 6, 8, and 10 h. WT is the wild-type biosensor, and J231XX-B003X represents different optimized combinations. (**F–H**), Dose-Response curves of three candidate combinations in shake flasks, F**–**H are the results for 6, 8, and 10 h. Values are mean ± s.d. (*n* = 3 biologically independent samples)

**Table 1.** Parameters of biosensor performance with standard error of mean.

| Sensor | Sensitivity | K<sub>d</sub> (μM) | Dynamic range (FI/OD<sub>600 nm</sub>) | Inducer range (μM) | R<sup>2</sup> |
| --- | --- | --- | --- | --- | --- |
| WT (6 h) | 0.37 (± 0.10) | MD | 0–318.5 | 10–1,000 | 0.9786 |
| 100-30 (6 h) | 0.52 (± 0.08) | 42.0 (± 17.1) | 6.9–1,172.4 | 0.1–1,000 | 0.9895 |
| 111-30 (6 h) | 0.82 (± 0.06) | 10.0 (± 1.0) | 42.3–1,988.9 | 0.1–1,000 | 0.9957 |
| 105-30 (6 h) | 0.58 (± 0.11) | 15.0 (± 6.0) | 107.3–2,035.2 | 0.1–1,000 | 0.9760 |
| fT-100-30 (6 h) | 0.59 (± 0.13) | 85.2 (± 45.0) | 9.0–306.4 | 3–1,000 | 0.9657 |
| fT-111-30 (6 h) | 0.77 (± 0.14) | 17.0 (± 4.1) | 40.8–599.9 | 1–500 | 0.9590 |
| fT-105-30 (6 h) | 0.83 (± 0.12) | 69.2 (± 15.0) | 153.9–402.2 | 3–1,000 | 0.9772 |
| WT (8 h) | 0.85 (± 0.16) | 8.7 (± 2.0) | 0–427.0 | 1–1,000 | 0.9905 |
| 100-30 (8 h) | 0.71 (± 0.07) | 12.6 (± 2.1) | 2.3–1,773.5 | 0.1–1,000 | 0.9917 |
| 111-30 (8 h) | 1.25 (± 0.09) | 6.8 (± 0.4) | 107.5–2,517.3 | 1–100 | 0.9972 |
| 105-30 (8 h) | 0.85 (± 0.10) | 9.2 (± 1.3) | 339.5–4,180.8 | 1–1,000 | 0.9905 |
| fT-100-30 (8 h) | 0.95 (± 0.18) | 77.0 (± 17.5) | 45.0–262.6 | 10–500 | 0.9651 |
| fT-111-30 (8 h) | 1.08 (± 0.12) | 26.2 (± 3.7) | 100.9–590.1 | 3–500 | 0.9762 |
| fT-105-30 (8 h) | 1.34 (± 0.19) | 45.0 (± 5.0) | 256.2–1,241.2 | 6–500 | 0.9772 |
| WT (10 h) | 0.59 (± 0.12) | 26.3 (± 12.3) | 0–581.5 | 1–1,000 | 0.9682 |
| 100-30 (10 h) | 0.73 (± 0.09) | 17.2 (± 3.3) | 2.9–2,327.0 | 0.1–1,000 | 0.9889 |
| 111-30 (10 h) | 0.74 (± 0.11) | 13.2 (± 3.0) | 78.5–4,434.4 | 0.1–1,000 | 0.9843 |
| 105-30 (10 h) | 0.62 (± 0.10) | 17.4 (± 5.7) | 348.0–5,203.0 | 0.1–1,000 | 0.9793 |
| fT-111-30 (10 h) | 1.33 (± 0.27) | 87.5 (± 11.4) | 169.1–2,366.2 | 10–500 | 0.9696 |
| fT-100-30 (10 h) | 1.47 (± 0.15) | 122.4 (± 9.7) | 69.3–1,150.7 | 10–500 | 0.9932 |
| fT-105-30 (10 h) | 1.14 (± 0.16) | 113.3 (± 17.0) | 412.3–4,323.0 | 10–1,000 | 0.9844 |

Sensitivity: Hill slope of fitted data. K<sub>d</sub>: concentration of SA to achieve half predicted maximal fluorescence intensity (FI)/OD<sub>600 nm</sub> value.
Dynamic range: minimal and maximal predicted FI/OD<sub>600 nm</sub> values. Inducer range: the range of SA which can be detected by the biosensor determined by experiments.
WT is the wild-type biosensor. 1XX-3X represents different optimized combinations. fT-1XX-3X indicates that the triple-plasmid transformant contains pSB1C3-1XX-3X (optimized biosensor), pA1a-P<sub>fic</sub>-*nahGH* (biodegradation) and pS8K-toxin/antitoxin (lethal circuit). All parameters were obtained by analysis of data measured in shake flasks. MD means meaningless data. Values are mean ± s.d. ( $n = 3$ biologically independent samples).

Due to limitations of oxygen and mass transfer in a 96-well plate, their functions were tested again in shake flasks. Compared with WT at 6 h (Fig. 1F and Table 1), the sensitivity of each promoter increased in the order of middle (J23111), low (J23105), and high (J23100) promoter intensity, which was consistent with the order of decreases of half-maximal activation concentration K_d_. Their dynamic range was extended, especially the maximum output, which changed from 318.5 to 2,035.2, while the order of the maximum outputs was opposite of the promoter intensities. Optimized sensors could detect 0.1 μM SA, which was two orders of magnitude lower than WT. Over time, the sensors became more sensitive, and the highest sensitivity was obtained at 8 h (1.27 of J23111-B0030, Fig 1G–1H and Table 1). However, the K_d_ values of J23111-B0030 and J23105-B0030 at 10 h were higher than at 6 h. Dynamic range was widened due to the continuous differential expression of mRFP, even though leaky expression became more significant. Except for J23111-B0030 and J23105-B0030 at 8 h, the detection range was 0.1–1,000 μM SA for each combination at all time periods. Meanwhile, the WT improved on some key parameters of the biosensor; for example, the maximum output of WT changed from 318.5 to 581.5, which still lagged behind the optimized groups.

### Stationary-phase biodegradation of salicylic acid

Stationary-phase promoters respond to starvation and cellular stress by transcribing downstream genes via the RNA polymerase containing the σ^S^ subunit (*16*). In the time dimension, they turn on the gene expression when strains grow to the stationary phase in rich media. We amplified five promoters from the *E. coli* BL21 (DE3) genome using the primers used by a previous study (*17*), and used mRFP as a reporter to characterize these promoters and select the best two. Three criteria were established: 1) the cell growth was not affected; 2) the start of transcription was strict, and the natural stationary-phase promoters were induced early in the late exponential phase (*17*); 3) it had a detectable output intensity. P_*katE*_ did not show any activity, while P_*osmY*_ and P_*csiE*_ turned on much earlier than the late exponential phase (>2 h). P_*bolA*_ and P_*fic*_ that met the above criteria were characterized with salicylate 5-hydroxylase (S5H), and the P_*bolA*_ was approximately 3-fold stronger than P_*fic*_. The growth-degradation curves at 1 mM SA were then plotted with the plasmid vector, pA1a. The engineered strains grew to a late exponential phase at about 8 h and SA began to be degraded after 6 h. The strain with P_*bolA*_ completely degraded SA in 16 h, and another one with P_*fic*_ within 12 h, which was opposite of the intensities characterized by mRFP. This is because a much stronger promoter could cause protein misfolding (*18*). As a result, we regarded P_*fic*_ as the optimal stationary-phase promoter for constructing the degradation circuit.

### Construction and characterization of lethal circuits

Several toxic proteins were selected to test their functions: CcdB, NucB, HokD, MazF, Gp2, RelK, and ProE. When the expression of toxic proteins was induced with Ara, strains containing HokD, MazF, RelK, ProE or Gp2 did not show any growth differences compared to the non-induced group. Unfortunately, positive transformants from CcdB were never obtained because the basal CcdB expression was sufficient to kill cells. To avoid this, intein was used to decrease the toxicity of intact CcdB, which was embedded in the host protein and could be autocatalytically excised during protein splicing before producing the mature protein (*19*). We determined that NucB, CcdB-42L, and CcdB-46V inhibited growth after 4 h of induction, while the lethal effect of CcdB-42L was better than that of 46V.

Lethal circuits were designed by two different concepts. The first one is based on a “gene converter” using NucB and CcdB-42L, respectively, where the CI repressor binds to the cI regulator and blocks the expression of downstream genes (*20*). Therefore, SA will activate the CI expression that suppresses the following toxic genes. The other was constructed by a pair of toxin/antitoxin (T/AT) proteins—CcdB-42L (toxin) and CcdA (antitoxin). CcdA and CcdB form a strong complex that prevents CcdB from interacting with the DNA gyrase (*21, 22*). When SA is present, the strain survives because of the expression of CcdA, whereas CcdB will cause cell death if SA is consumed. When all lethal circuits were compared in the cell growth curves, only the T/AT circuit showed a negative effect. Colony-forming unit (CFU) is a common method of characterizing the lethal or survival ratio; in the T/AT circuit, survival cells were 10^3^ times more than dead cells, while the others showed no difference at 8 h. After 4 h, the dead cells increased due to the accumulation of toxic proteins, and the T/AT circuit maintained the most powerful lethal ability, the survival ratio of which decreased to 10^−5^. According to the CFU, NucB was better able to induce cell death than CcdB-42L in a converter circuit. Therefore, we used the T/AT circuit as the lethal system in our engineered strain.

### Integration of functional modules and characterization of intelligent strains

To make the strain sequentially complete the sensing, degradation, and lethality, three modules were integrated into one strain and tested its functions. We first obtained a double-plasmid transformant containing degradation and lethality. The characteristics of the stationary-phase promoter did not change (Fig. 2A), i.e., the degradation started and ended at the same time as the single transformant. CFU for 20 h was measured owing to the accumulation of CcdA. The difference in growth curves did not appear, and the minimum survival ratio decreased to 10^−3^, which was similar to the data when the lethal circuit was separately tested at 8 h (Fig. 2A, 2B). The general trend of the survival ratio was that it maintained at a high level until SA was completely degraded and then decreased to a lower stage. In many cases, the SA concentration might be lower than 0.1 mM (*23*); therefore, we tested the strain at 0.1 mM SA. SA was degraded from 4 h onwards and was not detectable at 8 h (Fig. 2C). This was because the leaky expression of S5H would result in significant degradation of SA at low concentrations. The growth was inhibited, and the survival ratio was lower than at 1 mM SA (Fig. 2D). However, the survival ratio trend remained consistent with the former; the main reason for this phenomenon was the low expression of CcdA.

**Figure 2.**
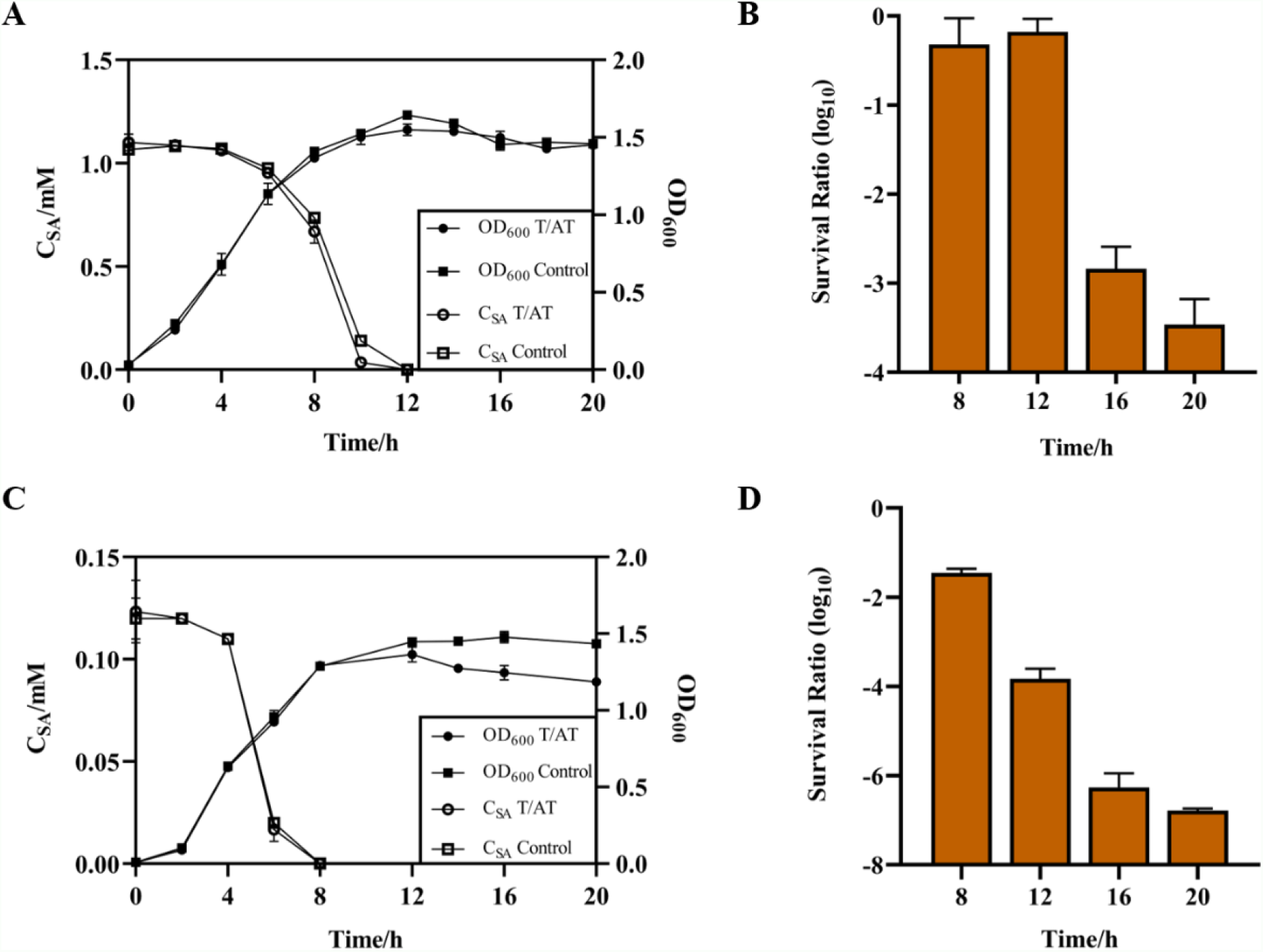
Characteristics of double-plasmid transformants containing biodegradation and lethal circuits. (**A–B**), Growth curves, degradation curves and survival ratios of double-plasmid transformants containing biodegradation and lethal circuit at 1 mM SA. Control is the double-plasmid transformant containing the biodegradation module and pS8K vector. (**C–D**), The above characterization at 0.1 mM SA. Values are mean ± s.d. (*n* = 3 biologically independent samples).

A biosensor was then introduced into the double-plasmid transformant. This triple-plasmid transformant responded to SA in the first few hours and began to degrade SA when the cells grew to the late exponential phase. Finally, a lethal circuit inhibited cell growth without SA. In the pre-selection at a 96-wells plate, the triple-plasmid combinations showed the same ability to detect SA as a single transformant (Fig. 3A–3C). Likewise, we scaled up this experiment to shake flasks and set up more SA concentrations (Fig. 3D–3F). The inducer range narrowed and K_d_ increased from 6 to 10 h. Therefore, we decided that 6 h was the optimal point to read the data of sensors (Table 1). The sensitivity, dynamic range, and inducer range of P_*fic*_-TAT-105-30 (triple-plasmid transformant containing pA1a-P_*fic*_-*nahGH*, pSB1C3-105-30, and pS8K-toxin/antitoxin) were the best of the three (P_*fic*_-TAT-100-30, P_*fic*_-TAT-111-30, P_*fic*_-TAT-105-30) although its K_d_ was slightly higher than P_*fic*_-TAT-111-30 at 6 h.

**Figure 3.**
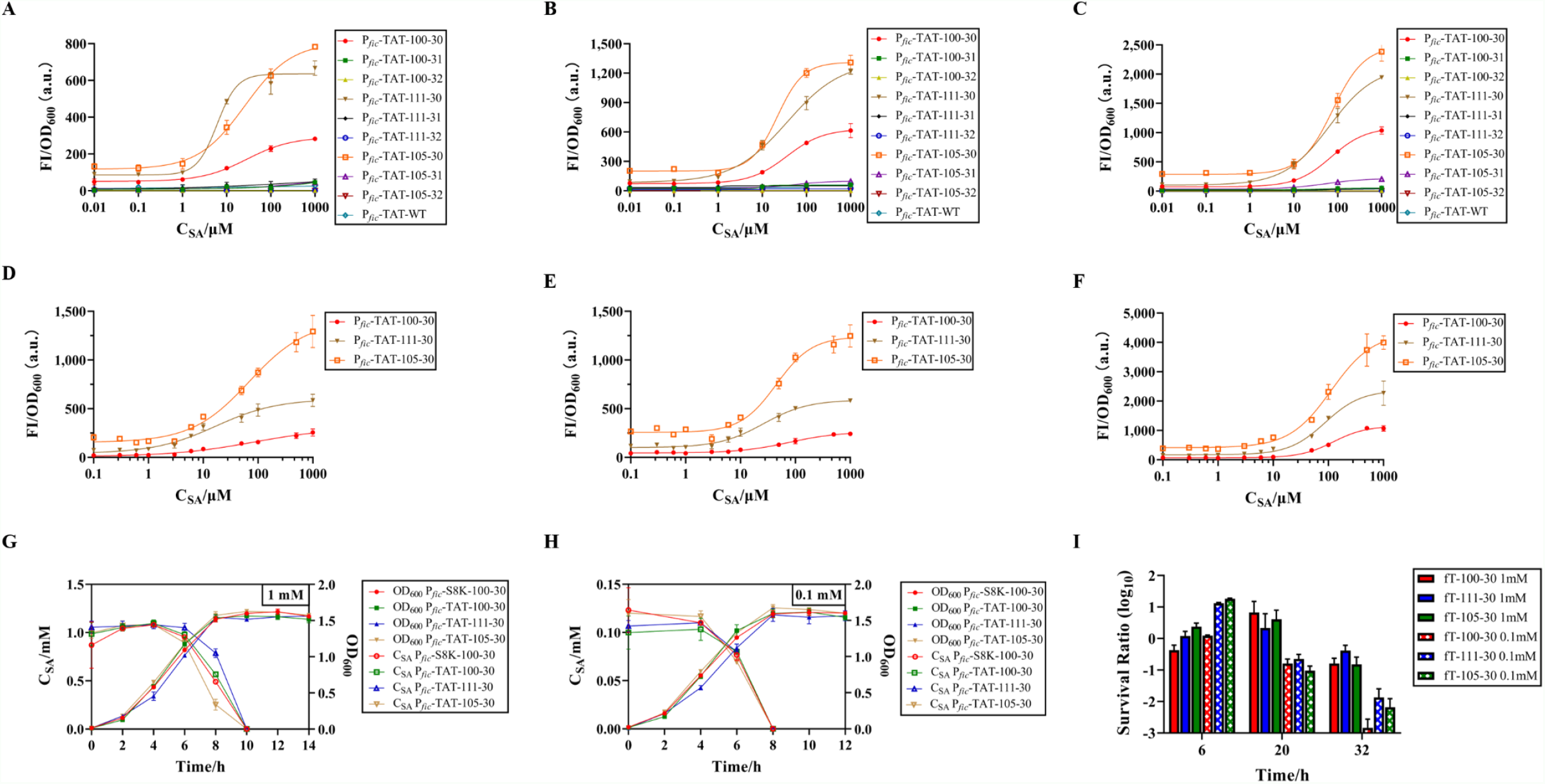
Characterization of triple-plasmid transformants containing the biosensors, biodegradation module, and lethal circuit. (**A–C**), Dose-Response curves of the triple-plasmid transformant with different optimized biosensors in 96-well plates, A–C are the results for 6, 8, and 10 h. P_*fic*_-TAT-1XX-3X/fT-1XX-3X indicates that the triple-plasmid transformant contains pSB1C3-1XX-3X (optimized biosensor), pA1a-P_*fic*_-*nahGH* (biodegradation module) and pS8K-toxin/antitoxin (lethal circuit), WT is the wild-type biosensor. (**D–F**), Dose-Response curves of the three best candidates in shake flasks. D–F are the results for 6, 8, and 10 h. (**G–I**), Growth curves, degradation curves, and survival ratios of the triple-plasmid transformant containing the biosensor, biodegradation module, and lethal circuit at 1 mM and 0.1 mM SA. P_*fic*_-S8K-100-30 is the triple-plasmid transformant containing the biosensor, biodegradation module, and pS8K vector as a control. Values are mean ± s.d. (*n* = 3 biologically independent samples).

Degradation and lethality were also tested at different SA concentrations (Fig. 3G–3I). SA of 1 mM was degraded faster than the double-plasmid transformant with a visible degradation from 4 to 6 h, but 0.1 mM SA was degraded the same as before. Unfortunately, no inhibition occurred in the growth curves under either concentration. The lethal effect also significantly decreased, with an approximately 10-fold difference of CFU in the survival group compared with the lethal group at 1 mM SA until 32 h. Even though all survival ratios of different groups at 0.1 mM SA decreased from 20 h, the minimum survival ratio at 32 h was only 2 orders of magnitude lower than those at 1 mM SA. Base on a t-test, there was no significant difference between the survival ratio of P_*fic*_-TAT-105-30 and P_*fic*_-TAT-111-30 at 0.1 mM SA for 32 h, which were only one order of magnitude higher than P_*fic*_-TAT-100-30. In summary, we regarded P_*fic*_-TAT-105-30 as our final engineered strain, even though its suicide system was not the most efficient.

## Discussion

We first optimized each of the three integrated modules. For detection, two common but effective methods were used to optimize our biosensor: 1) regulating the density of the receptor and 2) modulating the intensity of the reporter, which have been already applied in the valerolactam and caprolactam biosensor (*24*). Our optimized biosensor (J23111-B0030 at 6 h) performed an around 2-fold increase in sensitivity, a 6-fold increase in maximum output, and two orders of magnitude decrease in detection limits compared to WT. The former strategy could change the sensitivity and detection range, while the latter was used to increase the output signal (Fig. 1C– 1H, Table 1). Although an extremely weak promoter could cause high basal expression or low output signals, the strongest promoter did not create the best biosensor (*25*). Compared with our optimal SA biosensor, Lux- and GFP-based *Acinetobacter* exhibited narrower salicylate detection ranges which were 1–100 μM and 10–100 μM, respectively (*26*). The wild MarR-P_*marO*_ sensor in *E. coli* required 24 h as a response time, which was six times longer than ours (*27*). Using CmeR in *E. coli* could only detect 0.1–1 mM salicylate in 20–24 h (*28*). There were many other ways to optimize the biosensor, such as promoter and RBS engineering, replication origin engineering, regulator protein engineering and cascaded amplifiers (*25, 29–31*). We assessed how time effected biosensor properties and found that sufficient time was required to produce responding signals and reach the optimal state, but too much time limited the application of biosensors (*3*). Meanwhile, computer-assisted tuning approaches like deep learning and machine learning predict the performance of optimized biosensors (*32*).

As for the degradation module, we collected several common stationary-phase promoters and characterized them with mRFP, focusing on their initiation times and intensities. Some had an earlier start time of more than 2 hours, which required RT-qPCR to demonstrate whether it was a leaky expression or its own feature. As in metabolic regulation, stronger promoters did not result in faster degradation. Therefore, it is necessary to assemble a stationary-phase promoter with appropriate strength and induction time for a specific enzyme, which can be achieved by *de novo* synthesis and promoter engineering (*33, 34*).

In the lethal circuit, for an activator-based transcription factor, repressor CI or T/AT pair was used to control the function of the toxic protein. Many toxic proteins are difficult to be constructed into circuits due to their powerful functions, such as CcdB. Splitting these proteins apart can decrease the toxicity from leaked expression and maintain their original functions (*35*). However, selecting the optimal split site limits its application. To reduce the selection scope, existing databases and mathematical simulation tools can predict the effect of split sites (*36, 37*). The weaker lethality of the “gene converter” could be because the CcdB expression could not be inhibited by the survival signal (1 mM SA), and the intensity of the P_R_ promoter was not high enough. The performance of the T/AT circuit is closely related to the expression of antitoxin and the relative content of toxin and antitoxin (*38*).

Moreover, we coupled modules into one strain step-by-step, focusing on the effectiveness of chronological control and the integrity of module function. When degradation and the lethal circuit were first combined, the survival ratio trend changed with the SA concentration which could cause varied CcdA expression. A delay between completion of degradation and the lowest survival ratio occurred, which highlights the benefits of using proteases, riboswitch-integrase combined platform, or other means to accelerate switching the regulatory proteins on and off (*39–41*). For the integrated system containing all three modules, each module was disturbed at a different level. Due to the degradation of SA over time, the biosensor characteristics changed more than the single transformant. Undoubtedly, the response time of the biosensor must be modulated by altering the growth medium or adding a hydrolysis label (*42*). After detection, the start and end time of degradation at 1 mM SA were both 2 h earlier because the stationary-phase promoter was deeply influenced by growth stress (*12*), and three plasmids might bring an extra metabolic burden. Unfortunately, both the biosensor and the lethal circuit have deteriorated because it was difficult to balance the NahR expression in these two genetic circuits to avoid crosstalk of identical parts, or the metabolic burden affected the function of each module. Like metabolic engineering, computer-assisted design can accurately regulate the expression intensity of a specific protein or replace the same part with another functional regulatory element such as a riboswitch (*27, 41, 43*). Additionally, to make circuits genetically stable and not hinder the host, they can be introduced into highly-insulated genomic landing pads (*44*). The ADPWH_Nah combined detection and degradation, but the detection range was limited by the weak activity of the degradation cluster (*13*). Biodegradation and biosafety modules were integrated into the BL21 (DE3) AI-GOS strain. However, an exogenous inducer was needed to activate the killing circuit (*45*). This is the first time that three functional modules (biosensor, biodegradation, biosafety) were integrated into one strain and efficiently completed their corresponding tasks in chronological order without any exogenous inducers.

In conclusion, we designed an integrated engineered strain that can perform sensing, degradation, and lethality, in chronological order, without any exogenous inducers. This strain could respond to 3–1,000 μM SA within 6 h; when the strain reached to the late exponential phase, the stationary-phase promoter began transcription of *nahGH* to degrade SA. Finally, in the absence of SA, our engineered strain killed themselves to ensure biosafety. This work optimizes each module to make them more powerful and regulates the integrated system using logic gates and chronologically-controlled parts, which solves the challenge of decentralized and inefficient functional modules.

## Methods and Materials

### Strains, plasmids, chemicals, and growth conditions

All plasmid cloning and characterization of the engineered genetic circuits were performed in *E. coli* Top10. Four plasmid backbones with different copy numbers, pS8K (low copy number), pA1a (middle), pSB1C3 and J61002 (high), were used to construct plasmid derivatives. All strains were cultured in Luria-Bertani (LB) medium containing 10 g L^-1^ tryptone, 10 g L^-1^ NaCl, and 5 g L^-1^ yeast extract with appropriate antibiotics. Generally, the concentration of ampicillin, kanamycin, and chloramphenicol were 100, 50, and 25 mg L^-1^, respectively. However, the concentrations of antibiotics were halved in the experiment with the triple-plasmid transformant. The concentrations of arabinose (Ara) and salicylic acid (SA) were 1 mol L^-1^ and 100 mmol L^-1^, respectively. Antibiotics, Ara, and SA were dissolved in ddH_2_O and filtered using 0.22 μm filters (Sango Biotech., F513161-0001).

All engineered strains were first inoculated from individual colonies on LB solid plates to an appropriate volume of LB liquid medium, and cultured overnight at 37 °C with shaking (200 r.p.m.). For characterization, the seed cultures were then diluted 100-fold into a fresh LB liquid medium under the same culture conditions.

### Genetic circuits construction and transformation

Standard polymerase chain reaction (PCR) amplification and Gibson assembly were used to construct the genetic circuits and plasmids (Vazyme, C112). *De novo* synthesized genes were purchased from BGI, China. The plasmids were transformed into *E. coli* Top10 following standard protocols, and the resulting engineered strains were confirmed by Sanger sequencing (BGI).

### Characterization of biosensors

NahR (regulator), mRFP (reporter), and P_*sal*_ (cognate promoter of NahR) were used to design the biosensor. Constitutive promoters and ribosome binding sites (RBS) with different strengths were introduced into the circuit by PCR. The seed cultures were diluted into an LB liquid medium containing gradient concentrations of SA, after which 200 μL of diluted culture was added to 96-well plate for incubation to select the optimal combination of promoter and RBS (microporous plate oscillator, MBR-420FL, Taitec). Samples were obtained at 6, 8, and 10 h. The fluorescence intensity (FI) of mRFP was measured by Tecan Spark fluorometry (583 ± 10 nm for excitation, 607 ± 10 nm for emission, gain = 120). At the same time, the optical density (OD_600 nm_) was read to represent cell density. The medium background of FI and OD_600 nm_ was determined by blank wells within fresh LB liquid medium and subtracted from the experiment groups. Data were processed by GraphPad Prism, and the dose-response curve was fitted using the Sigmoidal, 4PL model. Candidates for the best combination were scaled up in a 250 mL flask containing 50 mL of medium. At least three experimental replicates were implemented for each experiment unless otherwise indicated.

### Characterization of biodegradation

Stationary-phase promoters, amplified from *E. coli* BL21 (DE3) by PCR (17), were designed to express the salicylate 5-hydroxylase (S5H). The strains with different stationary-phase promoters and *mrfp* were cultured in 96-well plates to compare their activities. FI and OD_600 nm_ were read at 0.5 h intervals until 12 h. For the best two, *mrfp* was replaced by *nahGH* and constructed in the pA1a, strains were scaled up in 250 mL flasks containing 100 mL LB and 1 mM SA. Samples were obtained every 2 hours until SA was completely consumed. The OD_600 nm_ was measured as described above and the concentration of SA was detected by high-performance liquid chromatography (HPLC) (Agilent Technologies 1200 series) with an Agilent Eclipse XDB-C18 column (5 μm, 4.6 × 150 mm). The HPLC parameters were as follows: flow rate 0.5 mL_•_min^-1^, flow phase 50% methanol and 50% deionized water with 0.1% (vol/vol) formic acid, column temperature 30°C, detection wavelength 298 nm, and stop time 15 min.

### Characterization of lethal systems

Several toxic proteins with different mechanisms were first compared first in 50 mL flasks containing 10 mL LB. OD_600 nm_ was measured by sampling every 2 h until 12 h, and incubation was induced with a final concentration of 10 mM Ara at 2 h. The split sites of CcdB were determined using the methods of the iGEM project (2019.igem.org/Team:DUT_China_B), except that 42L was used in a previous study (*19*). Two toxic proteins were used to design lethal circuits by two different concepts—CcdB and NucB for gene converter, respectively, and CcdB for toxin/antitoxin pair. The engineered strains were cultured under survival conditions (containing SA at a final concentration of 1 mM) and dead conditions (without SA). To characterize the lethal efficiency, growth curves and survival ratios were measured in 250 mL flasks containing 50 mL LB. Survival ratios were calculated based on CFU using the following formula:

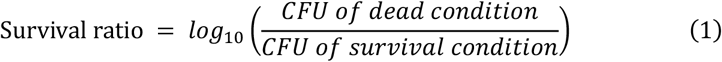

CFU is the number of single colonies on the agar plate, which is more accurate when this number is between 30 and 300. Samples were serially diluted to a proper concentration, and 200 μL of the diluent was spread on the LB solid plate, which was incubated upside down at 37°C overnight. To measure the expression of toxic protein in the ‘gene converter’ circuit at the ‘off’ state, *mrfp* was introduced into different circuit positions, and FI was measured as described above.

### Transformation and characterization of multiple-plasmid strains

New plasmids were introduced into the engineered strain that already contained one or two plasmids using standard electrotransformation protocols. For the double-plasmid transformant containing biodegradation and lethal system modules, it was cultured in a 250 mL flask containing 50 mL LB at a final concentration of 1 mM and 0.1 mM SA, respectively. Samples were obtained to measure the OD_600 nm_, concentrations of SA and survival ratio for 20 h. Characterization of triple-plasmid transformant was the same as the double one for 32 h, expect for an extra biosensor parameter, FI.

## Statistical analysis

Statistical analysis was performed by GraphPad Prism 8.0. Values are mean ± s.d. (*n* = 3 biologically independent samples). Significance analysis between the two data was conducted by t-test using SPSS. The dose-response curve was fitted using the Sigmoidal, 4PL model in GraphPad Prism 8.0.

## Author contributions

H. Liu, L. Zhang, and H. Tang outset and designed experiments. H. Liu and L. Zhang performed experiments. H. Liu, L. Zhang, H. Hu, W. Wang, and H. Tang analyzed the data. H. Tang and P. Xu received projects, contributed reagents and materials. H. Liu, L. Zhang, and H. Tang wrote the paper. All Authors importantly discussed and revised the manuscript. All Authors commented on the manuscript before submission. All authors read and approved the final manuscript.

## Data and materials availability

All data are available in the main text or the supplementary materials.

## References

1. L. Xiang, G. Li, L. Wen, C. Su, Y. Liu, H. Tang, J. Dai, Biodegradation of aromatic pollutants meets synthetic biology. Synth. Syst. Biotechnol. 6, 153–162 (2021).

2. Q. Gui, T. Lawson, S. Shan, L. Yan, Y. Liu, The application of whole cell-based biosensors for use in environmental analysis and in medical diagnostics. Sensors 17, 1623 (2017).

3. N. Ding, S. Zhou, Y. Deng, Transcription-factor-based biosensor engineering for applications in synthetic biology. ACS Synth. Biol. 10, 911–922 (2021).

4. H. Chong, C. B. Ching, Development of colorimetric-based whole-cell biosensor for organophosphorus compounds by engineering transcription regulator DmpR. ACS Synth. Biol. 5, 1290–1298 (2016).

5. L. Rucka, J. Nesvera, M. Patek, Biodegradation of phenol and its derivatives by engineered bacteria: current knowledge and perspectives. World J. Microbiol. Biotechnol. 33, 174 (2017).

6. S. Manna, J. Truong, M. C. Hammond, Guanidine biosensors enable comparison of cellular turn-on kinetics of riboswitch-based biosensor and reporter. ACS Synth. Biol. 10, 566–578 (2021).

7. H. Habe, T. Omori, Genetics of polycyclic aromatic hydrocarbon metabolism in diverse aerobic bacteria. Biosci. Biotechnol. Biochem. 67, 225–243 (2003).

8. G. Zhang, X. Yang, Z. Zhao, T. Xu, X. Jia, Artificial consortium of three E. coli BL21 strains with synergistic functional modules for complete phenanthrene degradation. ACS Synth. Biol. 11, 162–175 (2022).

9. D. Z. Yan, L. Q. Mao, C. Z. Li, J. Liu, Biodegradation of hexachlorobenzene by a constructed microbial consortium. World J. Microbiol. Biotechnol. 31, 371–377 (2015).

10. C. T. Chan, J. W. Lee, D. E. Cameron, C. J. Bashor, J. J. Collins, ‘Deadman’ and ‘Passcode’ microbial kill switches for bacterial containment. Nat. Chem. Biol. 12, 82–86 (2016).

11. J. W. Lee, C. T. Y. Chan, S. Slomovic, J. J. Collins, Next-generation biocontainment systems for engineered organisms. Nat. Chem. Biol. 14, 530–537 (2018).

12. J. Jaishankar, P. Srivastava, Molecular basis of stationary phase survival and applications. Front. Microbiol. 8, 2000 (2017).

13. Y. Sun, X. Zhao, D. Zhang, A. Ding, C. Chen, W. E. Huang, H. Zhang, New naphthalene whole-cell bioreporter for measuring and assessing naphthalene in polycyclic aromatic hydrocarbons contaminated site. Chemosphere 186, 510–518 (2017).

14. Y. Xue, P. Du, A. A. Ibrahim Shendi, B. Yu, Mercury bioremediation in aquatic environment by genetically modified bacteria with self-controlled biosecurity circuit. J. Clean. Prod. 337, (2022).

15. J. Z. Huang, M. A. Schell, In vivo interactions of the NahR transcriptional activator with its target sequences. Inducer-mediated changes resulting in transcription activation. J. Biol. Chem. 266, 10830–10838 (1991).

16. S. Lacour, P. Landini, σ^S^-dependent gene expression at the onset of stationary phase in Escherichia coli: function of σ^S^-dependent genes and identification of their promoter sequences. J. Bacteriol. 186, 7186–7195 (2004).

17. G. Miksch, F. Bettenworth, K. Friehs, E. Flaschel, A. Saalbach, T. Twellmann, T. W. Nattkemper, Libraries of synthetic stationary-phase and stress promoters as a tool for fine-tuning of expression of recombinant proteins in Escherichia coli. J. Biotechnol. 120, 25–37 (2005).

18. F. Baneyx, Recombinant protein expression in Escherichia coli. Curr. Opin. Biotech. 10, 411–421 (1999).

19. R. Lopez-Igual, J. Bernal-Bayard, A. Rodriguez-Paton, J. M. Ghigo, D. Mazel, Engineered toxin-intein antimicrobials can selectively target and kill antibiotic-resistant bacteria in mixed populations. Nat. Biotechnol. 37, 755–760 (2019).

20. B. Saltepe, E. S. Kehribar, S. S. Su Yirmibesoglu, U. O. Safak Seker, Cellular biosensors with engineered genetic circuits. ACS Sens. 3, 13–26 (2018).

21. A. Harms, D. E. Brodersen, N. Mitarai, K. Gerdes, Toxins, targets, and triggers: an overview of toxin-antitoxin biology. Mol. Cell 70, 768–784 (2018).

22. E. M. Bahassi, M. H. O’Dea, N. Allali, J. Messens, M. Gellert, M. Couturier, Interactions of CcdB with DNA gyrase. Inactivation of Gyra, poisoning of the gyrase-DNA complex, and the antidote action of CcdA. J. Biol. Chem. 274, 10936–10944 (1999).

23. R. G. Combarros, I. Rosas, A. G. Lavín, M. Rendueles, M. Díaz, Influence of biofilm on activated carbon on the adsorption and biodegradation of salicylic acid in wastewater. Water Air Soil Poll. 225, 1858 (2014).

24. M. G. Thompson, A. N. Pearson, J. F. Barajas, P. Cruz-Morales, N. Sedaghatian, Z. Costello, M. E. Garber, M. R. Incha, L. E. Valencia, E. E. K. Baidoo, H. C. Martin, A. Mukhopadhyay, J. D. Keasling, Identification, characterization, and application of a highly sensitive lactam biosensor from Pseudomonas putida. ACS Synth. Biol. 9, 53–62 (2020).

25. X. Wan, F. Volpetti, E. Petrova, C. French, S. J. Maerkl, B. Wang, Cascaded amplifying circuits enable ultrasensitive cellular sensors for toxic metals. Nat. Chem. Biol. 15, 540– 548 (2019).

26. W. E. Huang, H. Wang, H. Zheng, L. Huang, A. C. Singer, I. Thompson, A. S. Whiteley, Chromosomally located gene fusions constructed in Acinetobacter sp. ADP1 for the detection of salicylate. Environ. Microbiol. 7, 1339–1348 (2005).

27. Y. Zou, C. Li, R. Zhang, T. Jiang, N. Liu, J. Wang, X. Wang, Y. Yan, Exploring the tunability and dynamic properties of MarR-PmarO sensor system in Escherichia coli. ACS Synth. Biol. 10, 2076–2086 (2021).

28. M. A. Nasr, L. R. Timmins, V. J. J. Martin, D. H. Kwan, A versatile transcription factor biosensor system responsive to multiple aromatic and indole inducers. ACS Synth. Biol. 11, 1692–1698 (2022).

29. A. J. Meyer, T. H. Segall-Shapiro, E. Glassey, J. Zhang, C. A. Voigt, Escherichia coli “Marionette” strains with 12 highly optimized small-molecule sensors. Nat. Chem. Biol. 15, 196–204 (2019).

30. Y. Chen, J. M.L. Ho, D. L. Shis, C. Gupta, J. Long, D. S. Wagner, W. Ott, K. Josić, M. R. Bennett, Tuning the dynamic range of bacterial promoters regulated by ligand-inducible transcription factors. Nat. Commun. 9, 64 (2018).

31. N. Ding, Z. Yuan, X. Zhang, J. Chen, S. Zhou, Y. Deng, Programmable cross-ribosome-binding sites to fine-tune the dynamic range of transcription factor-based biosensor. Nucleic Acids Res. 48, 10602–10613 (2020).

32. P. Carbonell, T. Radivojevic, H. Garcia Martin, Opportunities at the intersection of aynthetic viology, machine learning, and automation. ACS Synth. Biol. 8, 1474–1477 (2019).

33. G. Miksch, F. Bettenworth, K. Friehs, E. Flaschel, The sequence upstream of the -10 consensus sequence modulates the strength and induction time of stationary-phase promoters in Escherichia coli. Appl. Microbiol. Biotechnol. 69, 312–320 (2005).

34. J. Jaishankar, P. Srivastava, Strong synthetic stationary phase promoter-based gene expression system for Escherichia coli. Plasmid 109, 102491 (2020).

35. D. D. Martin, M.-Q. Xu, T. C. Evans, Characterization of a naturally occurring trans-splicing intein from Synechocystis sp. PCC6803. Biochemistry 40, 1393–1402 (2001).

36. T. B. Dolberg, A. T. Meger, J. D. Boucher, W. K. Corcoran, E. E. Schauer, A. N. Prybutok, S. Raman, J. N. Leonard, Computation-guided optimization of split protein systems. Nat. Chem. Biol. 17, 531–539 (2021).

37. F. B. Perler, InBase: the Intein Database. Nucleic Acids Res. 30, 383–384 (2002).

38. F. Stirling, L. Bitzan, S. O’Keefe, E. Redfield, J. W.K. Oliver, J. Way, P. A. Silver, Rational design of evolutionarily stable microbial kill switches. Mol. Cell 68, 686–697 (2017).

39. J. Fernandez-Rodriguez, C. A. Voigt, Post-translational control of genetic circuits using Potyvirus proteases. Nucleic Acids Res. 44, 6493–6502 (2016).

40. K. L. Griffith, A. D. Grossman, Inducible protein degradation in Bacillus subtilis using heterologous peptide tags and adaptor proteins to target substrates to the protease ClpXP. Mol. Microbiol. 70, 1012–1025 (2008).

41. H. L. Pham, A. Wong, N. Chua, W. S. Teo, W. S. Yew, M. W. Chang, Engineering a riboswitch-based genetic platform for the self-directed evolution of acid-tolerant phenotypes. Nat. Commun. 8, 411 (2017).

42. X. F. Chen, X. X. Xia, S. Y. Lee, Z. G. Qian, Engineering tunable biosensors for monitoring putrescine in Escherichia coli. Biotechnol. Bioeng. 115, 1014–1027 (2018).

43. M. Koch, A. Pandi, O. Borkowski, A. C. Batista, J. L. Faulon, Custom-made transcriptional biosensors for metabolic engineering. Curr. Opin. Biotechnol. 59, 78–84 (2019).

44. Y. Park, A. Espah Borujeni, T. E. Gorochowski, J. Shin, C. A. Voigt, Precision design of stable genetic circuits carried in highly-insulated E. coli genomic landing pads. Mol. Syst. Biol. 16, e9584 (2020).

45. Q. Li, Y.-J. Wu, A fluorescent, genetically engineered microorganism that degrades organophosphates and commits suicide when required. Appl. Microbiol. Biot. 82, 749– 756 (2009).

